# Generation and characterization of *Sema6aΔcyt* conditional knockout mice

**DOI:** 10.1101/2023.11.06.565851

**Authors:** Marieke G. Verhagen, Eljo Y. van Battum, Marleen H. van den Munkhof, Suzanne Lemstra, Kati Rehberg, Aikaterini Koutourlou, Klara Jansen, Christiaan van der Meer, Nicky C. H. van Kronenburg, Youri Adolfs, R. Jeroen Pasterkamp

## Abstract

The axon guidance molecule Semaphorin-6A (SEMA6A) plays a key role during nervous system development. SEMA6A, classically known as a ligand for Plexin-A2 and -A4, is a transmembrane protein that can also elicit signaling via its intracellular domain *in vitro*. However, the physiological relevance of this ‘*revers*e’ signaling route is largely unknown. We generated a new transgenic mouse model, *Sema6aΔcyt^fl/fl^*, in which the cytosolic part of SEMA6A can be conditionally removed using *Cre*-recombination. Upon *Sema6aΔcyt* mutation, SEMA6A can only act as a ligand and *reverse* signaling is perturbed. Germline deletion of SEMA6A’s intracellular part results in developmental defects in axon pathfinding and neuron migration that partially phenocopy defects observed in full *Sema6a* knockout mice. These defects include disorganization of the anterior commissure, piriform cortex, lateral olfactory tract, thalamocortical and corticospinal white matter tracts, and defected neuron migration in the neocortex and cerebellum. Intriguingly, the hippocampal malformation described in *Sema6a* full knockout mice was not reproduced, suggesting a specific role for SEMA6A *forward* signaling in hippocampal development. Our results indicate that the intracellular domain of SEMA6A is essential for proper axon targeting and neuron migration, and provide the first proof of a SEMA6A *reverse* signaling pathway *in vivo*.

## INTRODUCTION

During nervous system development, newborn neurons and growing neurites find their way through a complex code of attractive and repulsive signals^1,2^. Since the 1990s a handful ‘axon guidance protein’ families have been identified, including Semaphorins and their Plexin receptors^3,4^. To orchestrate the generation of highly intricate brain networks, additional layers of signaling complexity of these relatively few proteins must be in place. Multiple ways of diversification of axon guidance protein signaling have been postulated, such as the formation of heterodimers or trimers^5^, protein binding in *cis*, or *reverse* signaling of receptor-ligand complexes^6,7^. However, specific *in vivo* roles for these additional protein functions are still largely elusive.

The transmembrane protein Semaphorin-6A (SEMA6A) plays a key role during nervous system development. SEMA6A has been shown to act as a ligand for Plexin-A2 and -A4 receptors inducing growth cone collapse and axon repulsion^8–11^. Interestingly, upon stimulation with Plexin-A2 *in vitro*, SEMA6A activates a reverse signaling pathway through its own intracellular domain also resulting in cytoskeleton remodeling^11^. In *Sema6a* full knockout mice (*Sema6a^-/-^*), in which both the *forward* and a potential *reverse* pathway are perturbed, several axonal and cellular phenotypes were observed throughout the central nervous system and retina^8,10,12–25^. Since most of the affected cell types express SEMA6A or co-express SEMA6A and Plexin-A2 themselves, it is impossible to rule out a potential role for impaired *reverse* signaling pathway in these phenotypes^14,17,22,26^.

The first hints for a role for Semaphorin *reverse* signaling during *in vivo* nervous system development were found in *Drosophila*, where Semaphorin-1a (Sema-1a, a SEMA6A orthologue) function in synapse formation was found to be dependent on its intracellular domain^27^. Similarly, in a genetic model where Plexin-a1 can only act as a ligand, Sema-1a regulates the fasciculation of axons in the visual system^28^. Moreover, it was shown that binding of Rho GTPase regulators at the intracellular domain of Sema-1a is essential for motor axon pathfinding in the spinal cord^29,30^. In mice, axons from SEMA6A-expressing On direction-selective ganglion cells (On DSGCs) in the retina are attracted by Plexin-A2 and -A4 ligands expressed in the medial terminal nucleus (MTN) of the accessory optic system^20^. In turn, axons from the MTN are attracted to the nucleus of the optic tract (NOT) in a similar way^24^. Lastly, in the chick spinal cord SEMA6A induces aggregation of boundary cap cells (BCCs) upon their encounter with Plexin-A1-expressing motor neuron axons, thereby preventing motor neuron soma mislocalization in the periphery^31^. Although these findings strongly suggest cell-autonomous functions for SEMA6A as a receptor, it is hard to differentiate between forward and reverse signaling routes and functions in an environment where both options are still open.

To functionally discriminate between the different signaling pathways of SEMA6A at axon guidance choice-points *in vivo*, we designed a conditional transgenic mouse model that enables selective deletion of the SEMA6A cytoplasmic domain using Cre/Lox-mediated recombination. This *Sema6aΔcyt^fl/fl^* mouse model permits the study of nervous system development in the absence of SEMA6A *reverse* signaling, whilst *forward* signaling remains attainable. Comparison of full *Sema6a* knockout mice with *Sema6aΔcyt^Δ/Δ^* mice, in which SEMA6A’s intracellular part is deleted from the germline, reveals similar developmental defects in the organization of the anterior commissure, piriform cortex, lateral olfactory tract, thalamocortical and corticospinal white matter tracts, and in cellular migration in the neocortex and cerebellum. However, a defect in dentate gyrus neuron positioning was not phenocopied in *Sema6aΔcyt^Δ/Δ^* mice. These results suggest a prominent role for SEMA6A *reverse* signaling pathways during neurodevelopment, thereby highlighting the diversification of Semaphorin function and challenging the general idea of SEMA6A acting as a ligand alone.

## RESULTS

### *Sema6aΔcyt* construct design

To perturb SEMA6A *reverse* signaling, we first designed a *Sema6aΔcyt* construct in which exon 19 of murine *Sema6a* was replaced by a truncated version lacking the sequence coding for its intracellular domain (Fig. 1A). Full-length *Sema6a* and *Sema6aΔcyt* cDNA sequences were placed under the control of the *pCAG* promotor sequence and were followed by *GFP*. To validate correct protein expression and localization, *pCAG-Sema6a-GFP* and *pCAG-Sema6aΔcyt-GFP* constructs were transfected into neuroblastoma (N2A) cells. Then, surface protein expression was verified by anti-SEMA6A antibody on non-permeabilized cells and compared to cytosolic expression using anti-GFP antibody after fixation and permeabilization (Fig. 1B). Both *pCAG-Sema6a-GFP* and *pCAG-Sema6aΔcyt-GFP* constructs were able to induce GFP expression. Moreover, we observed a similar GFP/SEMA6A intensity ratio (0.668 ± 0.106 (*Sema6a-GFP*) vs. 0.814 ± 0.056 (*Sema6aΔcyt-GFP*), p = 0.200, Fig. 1C), indicating correct localization of the truncated SEMA6AΔcyt protein into the cell membrane. We then checked whether our *pCAG-Sema6a-GFP* and *pCAG-Sema6aΔcyt-GFP* constructs indeed produced the full-length or truncated protein, respectively. Western blot analysis of membrane fractions of N2A cells using anti-SEMA6A antibody indeed revealed cell surface expression of a 1031 amino acid (aa) *wildtype* SEMA6A protein after *pCAG-Sema6a-GFP* transfection (± 160 kDa band), and a smaller 631 aa protein upon *pCAG-Sema6aΔcyt-GFP* transfection (± 120 kDa band, Fig. 1D left panel). Similar-sized proteins were found in these membrane fractions with the use of anti-GFP antibody (Fig. 1D right panel).

**Figure 1.**
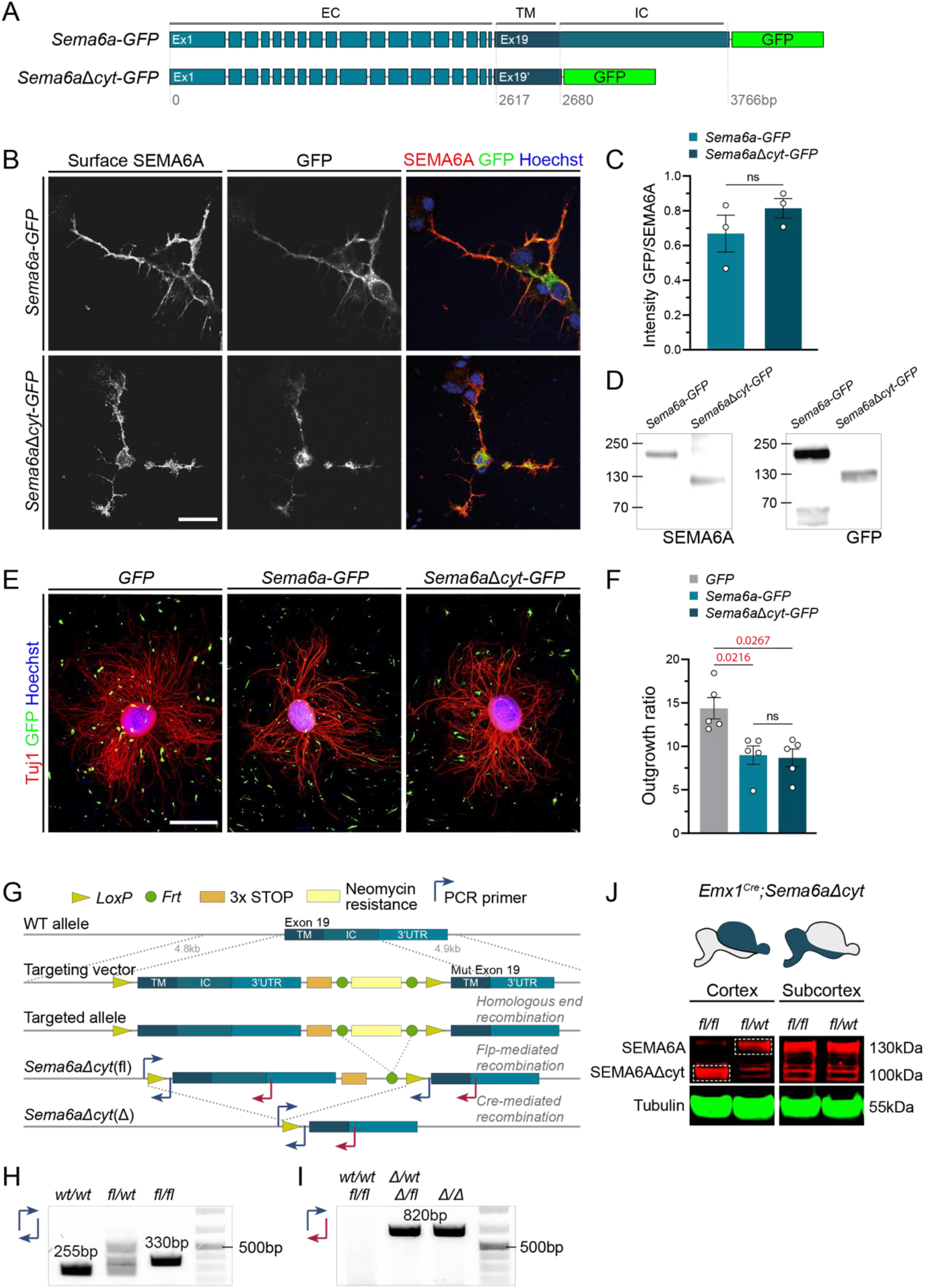
Design of *Sema6aΔcyt* construct and *in vitro* and *in vivo* validation experiments. **A)** Schematic representation of *Sema6a-GFP* and *Sem6aΔcyt-GFP* cDNA sequences. *Sem6aΔcyt-GFP* contains a truncated exon 19 (Ex19’), in which the sequence coding for the SEMA6A intracellular domain was deleted. **B)** Immunocytochemistry of surface SEMA6A expression (before permeabilization) and cytosolic GFP expression (after permeabilization) on N2A cells transfected with *pCAG-Sema6a-GFP* and *pCAG-Sem6aΔcyt-GFP* plasmids. **C)** Quantification of cell surface expression of SEMA6AΔcyt-GFP and SEMA6A-GFP in N2A cells as in (B). Values are calculated as the ratio between cytosolic GFP and surface SEMA6A staining intensity, counted from 103 (*Sema6a-GFP*) and 124 cells (*Sem6aΔcyt-GFP*) averaged from N=3 experiments, Mann-Whitney U test, p = 0.200, not significant). **D)** Western blot analysis of SEMA6A-GFP and SEMA6AΔcyt-GFP overexpressed in N2A cells. Cell lysates show SEMA6A-GFP peptide (160 kDa) and a truncated SEMA6AΔcyt-GFP peptide (120 kDa) using goat-anti-SEMA6A (left panel) and rabbit-anti-GFP (right panel) antibodies. **E)** *Wildtype* primary cortical explants after 48 h of culture on a NIH3T3 cell monolayer transfected with *pCAG-GFP* control, *pCAG-Sema6a-GFP*, or *pCAG-Sema6aΔcyt-GFP* plasmids. **F)** Quantification of explant outgrowth on N2A cells transfected with *pCAG-GFP*, *pCAG-Sema6a-GFP*, or *pCAG-Sema6aΔcyt-GFP* plasmids as in (E). Values were calculated as the ratio of total outgrowth size to explant core size, averaged from 53 explants (*GFP*), 72 explants (*Sema6a*) and 71 explants (*Sema6aΔcyt*) from N=5 experiments. One-way ANOVA with Kruskal-Wallis multiple comparison test. No significant change was detected between *Sema6aΔcyt* and *Sema6a*, p > 0.999. **G)** Design of targeting vector and BAC-mediated homologous recombination to generate the conditional *Sema6aΔcyt* mouse model. The *wildtype* full-length exon 19 was flanked with *LoxP* sites and a neomycin resistance between *Frt*-sites, and a shortened exon 19 (missing the part coding for the intracellular domain, but keeping the 3’UTR) was placed behind this. Together with two homologous arms (4.8 kb on 5’ side, and 4.9 kb on 3’ side) this combination was placed in a targeting vector, which was used to electroporate ES cells, and generate *Sema6aΔcyt* founders. *Flp*-mediated recombination resulted in excision of the neomycin cassette. *Cre*-mediated recombination results in either conditional (*Sema6aΔcyt(fl)*), or germline (*Sema6aΔcyt(Δ)*) removal of the wildtype exon 19, replacing it with the truncated version. **H)** Example of PCR analysis using primers flanking *LoxP* sites to determine *Sema6aΔcyt^fl^*genotype. PCR primer pair is indicated in (G) at the corresponding annealing location, and can be found in Table 3. **I)** Example of PCR analysis using primers located just before the *LoxP* site and just inside the 3’UTR, to determine *Sema6aΔcyt^Δ^* genotype. PCR primer pair is indicated in (G) at the corresponding annealing location, and can be found in Table 3. **J)** Validation of conditional *Sema6aΔcyt* mutation and truncated SEMA6A expression in postnatal *Emx1^Cre^;Sema6aΔcyt^fl/fl^* and *Emx1^Cre^;Sema6aΔcyt^fl/wt^*control mice. Schematic mouse brains indicate dissected cortical and subcortical samples in blue. Membrane fractions of *Emx1^Cre^;Sema6aΔcyt^fl/fl^*(and *Emx1^Cre^;Sema6aΔcyt^fl/wt^* to a lesser extend) cortical tissue show the expression of SEMA6AΔcyt protein (97 kDa). Membrane fractions of *Emx1^Cre^;Sema6aΔcyt^fl/wt^* cortices and both *Emx1^Cre^;Sema6aΔcyt^fl/fl^* and *Emx1^Cre^;Sema6aΔcyt^fl/wt^* subcortical samples show full length SEMA6A (130 kDa) expression. Scale bars 20 µm (B) and 500 µm (D). All data are represented as mean ± SEM Abbreviations: Bp, base pair, kb, kilobase; EC, extracellular; TM, transmembrane; IC, intracellular; WT, *wildtype*.

Next, we determined whether the SEMA6AΔcyt protein could still elicit *forward* signaling through its extracellular domain. To this end, cortical explants were placed on NIH3T3 cells transiently transfected with *pCAG-Sema6a-GFP, pCAG-Sema6Δcyt-GFP*, or *pCAG-GFP* control constructs, and assessed for outgrowth after 48 hours of culture (Fig. 1E). Compared to cortical explants grown on NIH3T3 cells expressing only GFP, explants showed reduced axon outgrowth when cultured on cells expressing full-length SEMA6A (14.37 ± 1.227 (GFP) vs. 8.992 ± 1.066 (SEMA6A), p = 0.0216) or SEMA6AΔcyt (14.37 ± 1.227 (GFP) vs. 8.671 ± 1.021 (SEMA6AΔcyt) p = 0.0267) (Fig. 1F). Together, these results indicate normal processing, expression and ligand function of the SEMA6AΔcyt protein.

### Generation of the *Sema6aΔcyt* mouse model

In order to study the function of SEMA6A *reverse* signaling *in vivo*, we generated a new transgenic mouse model, *Sema6aΔcyt^fl^*, in which the SEMA6A intracellular domain can be conditionally removed using *Cre*-recombination. We assembled a construct containing *wildtype Sema6a* exon 19 including the STOP codon and an Frt-flanked neomycin cassette inserted at the 5’ end. This entire sequence was flanked by *LoxP* sites and followed by a truncated *Sema6a* version of exon 19 lacking the coding part for the SEMA6A cytoplasmic domain (Fig. 1G). The vector was completed with 4.8 (5’-side) and 4.9 kb (3’-side) homologous arms at both ends of the inserted combination, and cloned into a pcDNA3.1 vector. The *Sema6aΔcyt* targeting vector was electroporated into ES cells (at Biocytogen). Neomycin (G418) resistant ES cells were selected and checked for correct integration of the construct by PCR and Southern blotting (data not shown). Transgenic ES cells were subsequently microinjected into pseudo-pregnant females (BALB/c mice) to generate chimeras and bred with *C57BL/6* mice to assess germline transmission resulting in F1 *Sema6aΔcyt^fl/wt^* mice. Three correctly targeted F1 founder mice were identified and maintained on a *C57BL/6* background. The Neomycin cassette was removed by Flp-mediated recombination (at Biocytogen). Subsequently, *Sema6aΔcyt^flfl^* mice were crossed with *Ella-Cre^Cre/wt^* mice^32^, after which offspring was selected for germline deletion of the SEMA6A intracellular domain, and crossed further to obtain *Sema6aΔcyt* null mice (*Sema6aΔcyt^Δ/Δ^*).

The presence of *loxP* sites in *Sema6aΔcyt^fl^* mice and germline deletion in *Sema6aΔcyt^Δ^* mice was confirmed by PCR analysis on genomic DNA derived from earclips. *LoxP* PCR resulted in a 255 basepair (bp) *wildtype* fragment and 330 bp mutant fragment (Fig. 1G,H). Germline deletion PCR resulted in a 820 bp mutant fragment, whereas amplification of a *wildtype* or *loxP*-flanked fragment was obstructed due to extended length (Fig. 1G,I).

Next, we validated that *Cre* recombination in *Sema6aΔcyt^fl/fl^*mice resulted in conditional deletion of the SEMA6A intracellular domain and that the resulting SEMA6AΔcyt protein remained correctly expressed at the cell membrane. Using *Emx1^Cre/wt^*mice^33^, which express *Cre* under control of the *Emx1* promotor in the cerebral cortex, we generated *Emx1^Cre/wt^;Sema6aΔcyt^fl/fl^*mice and performed western blot analysis on cortical and subcortical membrane fractions. Indeed, cortical membrane fractions of *Emx1^Cre/wt^;Sema6aΔcyt^fl/fl^*mice showed a truncated SEMA6AΔcyt protein of approximately 97 kDa, whereas *Emx1^Cre/wt^;Sema6aΔcyt^wt/wt^* mice displayed full-length SEMA6A protein of approximately 130 kDa (Fig. 1J). In subcortical membrane fractions, where no excision had occurred, full-length SEMA6A protein was detected in both *Emx1^Cre/wt^;Sema6aΔcyt^fl/fl^* and *Emx1^Cre/wt^;Sema6aΔcyt^wt/wt^* littermates (Fig. 1J). These results confirm correct excision of the SEMA6A intracellular domain and localization of the subsequent SEMA6AΔcyt protein at the cell membrane *in vivo*.

### Neurodevelopmental phenotypes in *Sema6aΔcyt* mice

It has been established that deletion of SEMA6A causes a broad range of neuroanatomical defects^8,10,12–25^. In order to disentangle the involvement of SEMA6A *reverse* and *forward* signaling in these developmental phenotypes, we assessed several of the anatomical phenotypes previously reported in *Sema6a^-/-^* mice in the newly generated *Sema6aΔcyt^Δ/Δ^*mice.

#### Defects in axon pathfinding

When new-born neurons reach their final location in the brain, their axons are guided towards local or long-distance targets to establish functional neural networks. Several defects in axonal pathfinding have been attributed to perturbed SEMA6A signaling, including misrouting of anterior commissural, thalamocortical and corticospinal projections^12,15–17,22^. We visualized and assessed these axonal tracts with L1CAM immunostaining in *wildtype, Sema6a^-/-^* and *Sema6aΔcyt^Δ/Δ^* mice.

First, axons of the posterior limb of the anterior commissure (pAC) normally cross the midline and establish interhemispheric connections. The pAC comprises axons of the agranular insular cortex, orbitofrontal cortices and some of the cingulate cortices^34^. We observed that in *wildtype* mice at postnatal day 0 (P0) pAC axons indeed cross the midline, whereas in *Sema6a^-/-^* and *Sema6aΔcyt^Δ/Δ^* mice these axons failed to project towards the midline and instead grew ventromedially alongside the piriform cortex (Fig. 2A,C). Moreover, in this same prefrontal region, DAPI nuclear staining highlighted excessive folding of the piriform cortex in *Sema6a^-/-^* as well as in *Sema6aΔcyt^Δ/Δ^* mice, which is evident by an enlarged curvature in the dense cell layer that comprises cortical layer 2 (LII) (Fig. 2B,C). Correspondingly, L1CAM^+^ axons from the lateral olfactory tract (LOT), that are normally located superficially at the piriform cortex in *wildtype* mice, were bundled together and extended more internally in the cortex of *Sema6a^-/-^* and *Sema6aΔcyt^Δ/Δ^* mice (Fig. 2A,C).

**Figure 2.**
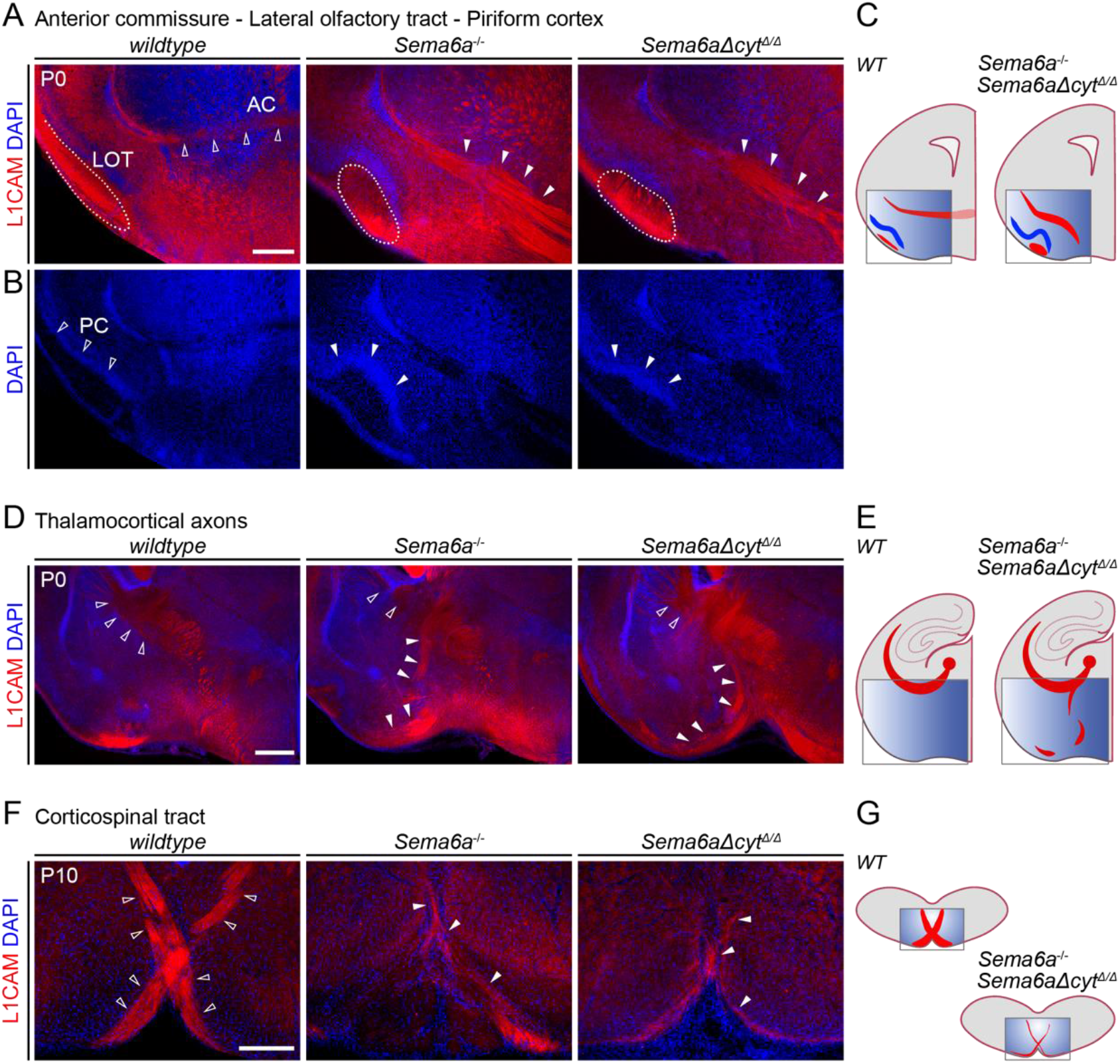
*Sema6aΔcyt^Δ/Δ^* mutants display similar axonal pathfinding defects as *Sema6a^-/-^* full knockout mice. **A,B,D,F**) Immunohistochemistry of coronal sections of P0 mouse brains from *C57Bl6/J* wildtype, *Sema6a^-/-^* and *Sema6aΔcyt^Δ/Δ^* mice. L1CAM staining reveals axonal bundles and DAPI shows cell nuclei. **A**) Axons running via the posterior limb of the anterior commissure (AC) cross the midline (*wildtype*, open arrowheads) below the telencephalic ventricles, are misplaced in both *Sema6a^-/^* and *Sema6aΔcyt^Δ/Δ^* mice. There, they fail to cross the midline and instead, run ventrally towards the hypothalamus (*Sema6a^-/^* and *Sema6aΔcyt^Δ/Δ^*, white arrowheads). The axons of the lateral olfactory tract (LOT, *wildtype*, dotted area), that normally form a superficial bundle along the piriform cortex, cluster together and form a thick bundle in *Sema6a^-/^* and *Sema6aΔcyt^Δ/Δ^* mice (dotted areas). **B**) The pyramidal cells of cortical layer 2 (here visualized by DAPI staining) forming the light curvature of the piriform cortex (PC, *wildtype*, open arrowheads), are folding much deeper in both *Sema6a^-/-^* and *Sema6aΔcyt^Δ/Δ^* mice (white arrowheads). **D**) Thalamocortical axons coming from the thalamus gather in the internal capsule (open arrowheads) to reach the cortex, however in both *Sema6a^-/-^* and *Sema6aΔcyt^Δ/Δ^* mice a subset is misprojecting ventrally (white arrowheads) running towards the surface of the ventral telencephalon. **F**) Axons of the cortical spinal tract cross the midline at the the pyramidal decussation (*wildtype*, open arrowheads). Hypoplasia of these corticospinal tracts is displayed by both *Sema6a^-/-^*and *Sema6aΔcyt^Δ/Δ^* mice (white arrowheads). Scale bars 500 µm (A, D), 200 µm (F). **C,E,G**) Schematic representation of axonal defects in *Sema6a^-/^* and *Sema6aΔcyt^Δ/Δ^* mice compared to *wildtype* (*WT*).

SEMA6A is also known to be involved in the guidance of thalamocortical axons (TCA)^12,16,22^. In *wildtype* conditions, thalamic axons have started to project through the internal capsule to reach the neocortex at P0 (Fig. 2D,E). Although some TCAs still projected towards the internal capsule in *Sema6a^-/-^* and *Sema6aΔcyt^Δ/Δ^* mice, both models displayed TCA bundles that deviate ventrally (Fig. 2C,D).

Lastly, defective axon pathfinding of the corticospinal tract (CST) was found in *Sema6a^-/-^* mice^15,23^. At the border between the medulla and spinal cord, pyramidal tracts normally project dorsally and cross the midline to form the pyramidal decussation and to enter the dorsal funiculus. As observed in *Sema6a^-/-^* mice, *Sema6aΔcyt^Δ/Δ^* CST axons showed severe hypoplasia at the level of the pyramidal decussation at P10 (Fig. 2F,G). Although a few remaining corticospinal axons still crossed the midline, the size of this bundle was markedly reduced and, in some instances, remained visible as a tight bundle at the ventral surface.

Taken together, we found that multiple of the axonal pathfinding defects described in *Sema6a^-/-^*mice were recapitulated in *Sema6aΔcyt^Δ/Δ^* mice, which suggests a crucial role for SEMA6A *reverse* signaling in the guidance of these projection paths.

#### Neuronal migration defects in the neocortex and cerebellum

Axon guidance proteins are known to not only guide axonal projections towards their correct targets, but also to regulate the migration of new-born neurons. For example, SEMA6A plays a key role in the migration of pyramidal neurons in the cerebral cortex and granule cells in the cerebellum^8,13,17,19,25^.

In the developing cortex, newly generated pyramidal neurons migrate radially in an inside-out pattern from the ventricular zone towards the outer cortex. The last-born neurons finish their migration in the uppermost cortical layers just below the marginal zone, which is a cell-sparse area that remains devoid of pyramidal neurons and will become cortical layer 1 (LI) after birth^35^. In the absence of SEMA6A signaling, however, these upper layer neurons overmigrate and invade LI^17,25^. The normally distinct border between the cell-sparse LI and the adjacent cell-dense LII is disturbed and neurons clustered together in the most superficial part of the cortex. This defect became more severe towards more lateral cortical regions (data not shown). We determined the presence of this cortical phenotype with NeuN immunostaining at P10. In both *Sema6a^-/-^* and *Sema6aΔcyt^Δ/Δ^* cortices, invasion and clustering of neurons into LI was detected (Fig. 3A,B). Mislocalization and clustering of Satb2^+^ upper layer neurons into LI was already visible at P4. No further laminar disturbances were detected by staining for markers labelling deeper layers of the cortex, such as Tbr1^+^ and Ctip2^+^, or LI (Reelin) (Fig. 3C). These data support previous findings showing that the mispositioned neurons have an upper layer identity^25^.

**Figure 3.**
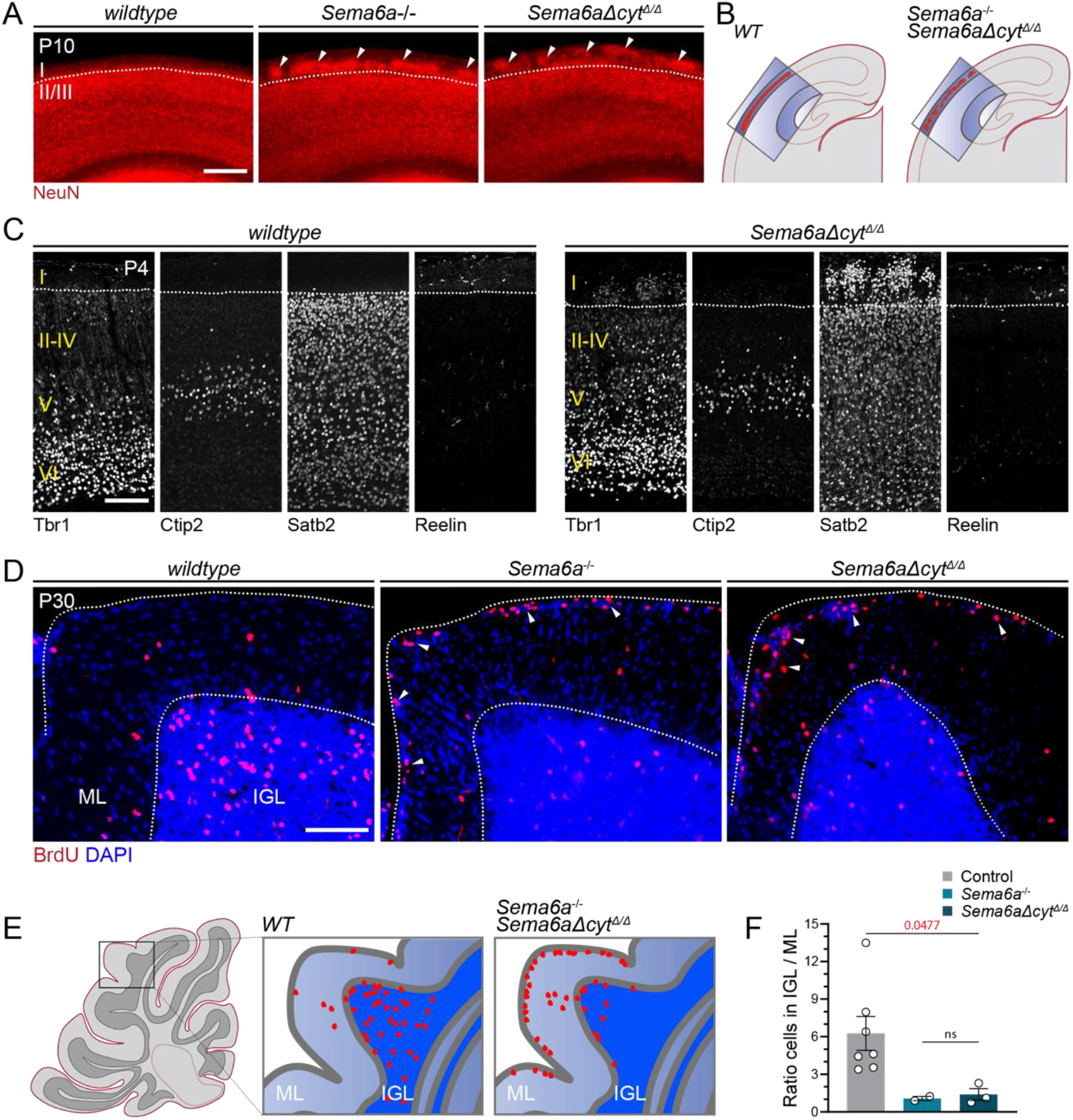
*Sema6aΔcyt^Δ/Δ^* mutants display similar migration defects in the neocortex and cerebellum like Sema6a^-/-^ knockout mice. **A)** Immunohistochemistry of sections of P10 *wildtype*, *Sema6a^-/-^* and *Sema6aΔcyt^Δ/Δ^* mice cortex, stained with neuronal marker NeuN. In the wildtype a smooth border between cortical layer I and layer II/III can be observed (dotted line). Both *Sema6a^-/-^*and *Sema6aΔcyt^Δ/Δ^* mice display overmigration of cortical neurons accumulating in clusters inside layer I (white arrowheads). **B)** Schematic representation of cortical migration defect described in A). **C)** Immunohistochemistry of cortical sections of P4 *wildtype*, *Sema6a^-/-^* and *Sema6aΔcyt^Δ/Δ^* mice, stained with markers for neurons residing in different cortical layers (Tbr1, cortical layer VI and V; Ctip2, cortical layer V; Satb2, cortical layer II-VI) and Cajal-Retzius cells (Reelin, layer I). **D)** Sagittal cerebellar sections of P30 *wildtype*, *Sema6a^-/-^*and *Sema6aΔcyt^Δ/Δ^* mice that were injected with BrdU at P15 to label a subset of cerebellar granule neurons. These neurons migrate through the molecular layer (ML) into the internal granule layer (IGL) of the cerebellum. In *Sema6a^-/-^*and *Sema6aΔcyt^Δ/Δ^* mice a large portion of BrdU-labeled cells stopped migration prematurely in the ML (white arrowheads). **E)** Schematic representation of cerebellar granule cell migration defect in *Sema6a^-/-^* and *Sema6aΔcyt^Δ/Δ^* mice, as in D). **F)** Quantification of BrdU-positive cerebellar granule cell migration towards the internal granule layer (IGL). A ratio was calculated between the amount of BrdU^+^ cells in the IGL and the molecular layer (ML). Ratios of 2 *C57Bl/6J*, 2 *Sema6a^+/-^* and 3 *Sema6aΔcyt^Δ/wt^* animals were pooled and serve as control condition to compare to N = 2 *Sema6a^-/-^* and N = 3 *Sema6aΔcyt^Δ/Δ^* mice. Kruskal-Wallis test with multiple comparison. Data is represented as mean ± SEM.

SEMA6A signaling also guides migrating neurons in the cerebellum, where it’s transiently expressed postnatally in migrating granule cells^13^. In the absence of SEMA6A, cerebellar granule cells, which normally switch from tangential migration in the external granule layer (EGL) to inwards radial migration in the molecular layer (ML) in order to reach the internal granule cell layer (IGL), stall and remain situated in the ML^8,13,19^. We verified these findings in *Sema6a^-/-^*and *Sema6aΔcyt^Δ/Δ^* mice by injecting bromodeoxyuridine (BrdU) intraperitoneally at P15, thereby targeting late-born cerebellar granule cells^19^, and examining their final positioning at P30. Indeed, whereas most of the BrdU-labelled cells managed to reach the IGL in *wildtype* mice, in *Sema6a^-/-^* and *Sema6aΔcyt^Δ/Δ^* mice BrdU-labelled granule cells halted their migration in the ML and were hardly detectable in the IGL (Fig. 3D,E). Accordingly, the ratio of BrdU-labelled granule cells in the IGL compared to the ML showed a five-fold decrease in *Sema6aΔcyt^Δ/Δ^* mice (Fig. 3F; 6.264 ± 1.354 (control) vs. 1.388 ± 0.463 (*Sema6aΔcyt^Δ/Δ^*), p = 0.0477). Due to the low number of animals available, no significant differences could be calculated between control and *Sema6a^-/-^* animals. However, the ratio of cerebellar granule cells that had migrated into the IGL of *Sema6a^-/-^* mice (1.069 ± 0.177, N = 2) was comparable to that found in *Sema6aΔcyt^Δ/Δ^* animals. These findings indicate that SEMA6A *reverse* signaling may be required for the proper radial migration of cortical upper layer neurons and cerebellar granule cells.

*Hippocampal defects are not phenocopied in Sema6aΔcyt^Δ/Δ^* mice

*Sema6a^-/-^* mice display a subtle malformation in the granule layer of the infrapyramidal blade of the dentate gyrus^17^. To assess hippocampal morphology in *Sema6a^-/-^* mice and our *Sema6aΔcyt* mouse model, we used Prox1 immunostaining to visualize post-mitotic dentate gyrus granule cells. Misplaced Prox1^+^ cells indeed cause a local broadening of the dentate gyrus in *Sema6a^-/-^* mice (Fig. 4). However, this defect was not observed in *Sema6aΔcyt^Δ/Δ^* mice, where the infrapyramidal blade was similar to *wildtype*. Further analysis of dentate gyrus morphology along the entire rostral-to-caudal axis of the hippocampus did not reveal any changes in *Sema6aΔcyt^Δ/Δ^* mice (Fig. 4D), indicating that development of the infrapyramidal blade of the dentate gyrus does not rely on the SEMA6A intracellular domain.

**Figure 4.**
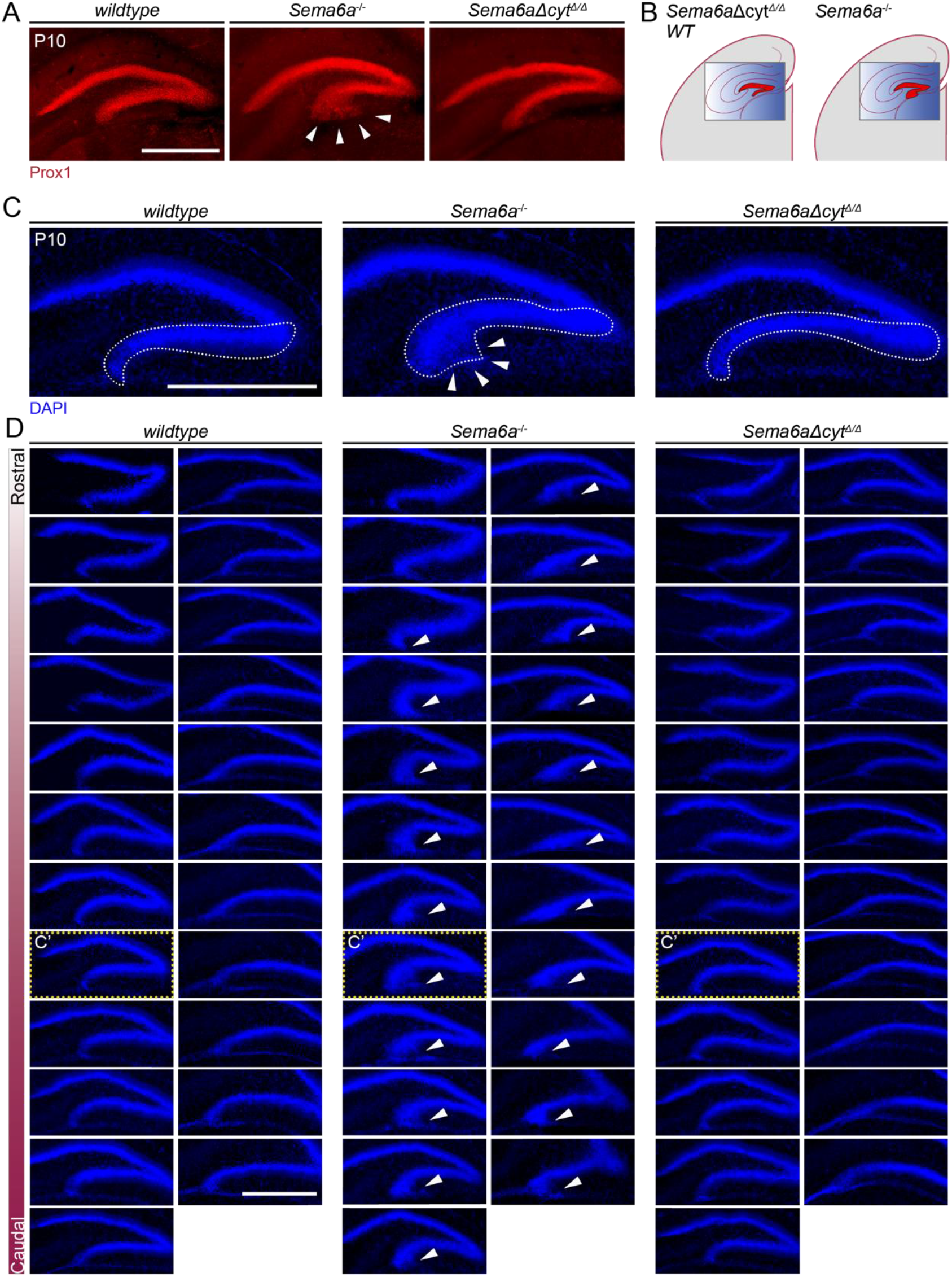
Malformation of the dentate gyrus is not recapitulated in *Sema6aΔcyt^Δ/Δ^* mice. **A)** Prox1 immunostaining labeling granule cells within the dentate gyrus of the hippocampus in *wildtype, Sema6a^-/-^* and *Sema6aΔcyt^Δ/Δ^* mice at P10. In *Sema6a^-/-^* mice, misplaced granule cells cause a broadened phenotype of the infrapyramidal blade of the dentate gyrus (white arrowheads), whereas *Sema6aΔcyt^Δ/Δ^* mice show a normal dentate gyrus morphology. **B)** Schematic representation of hippocampal phenotypes as in (A). **C)** DAPI staining at P10 showing morphology of the infrapyramidal blade (dotted lines) and a broadened dentate gyrus phenotype (white arrowheads) in *Sema6a^-/-^* mice. **D)** The dentate gyrus defect in *Sema6a^-/-^* mice is present along the rostral-to-caudal axis of the hippocampus. Panels depicted in (C) are indicated with C’. Scale bars 500 µm.

## DISCUSSION

The vertebrate axon guidance protein SEMA6A has the potential to elicit both *forward* signaling (as a ligand for Plexin-As) and *reverse* signaling (as receptor for Plexin-As) events. So far, functional evidence for the SEMA6A *reverse* pathway is supported by *in vitro* experiments where, depending on the cellular context, Plexin-A2 ectodomain stimulation induced collapse of SEMA6A-expressing cells^11^. Here, we present a novel mouse model in which the cytoplasmic domain of SEMA6A can be conditionally removed (*Sema6aΔcyt^fl^*) and provide the first evidence that *reverse* signaling of SEMA6A is essential for proper nervous system development *in vivo*.

Crosses of the *Sema6aΔcyt^fl/fl^* mice with *Emx1^Cre^*mice confirmed membrane expression of a truncated protein in the forebrain. Both *Emx1^Cre^;Sema6aΔcyt^fl/fl^* and *Sema6aΔcyt^Δ/Δ^* mice are viable and fertile, and display neurodevelopmental defects. A systematic comparison of *Sema6a^-/-^* full knockout mice and our novel *Sema6aΔcyt^Δ/Δ^* mice showed a striking overlap between phenotypes. Both *Sema6a^-/-^* and *Sema6aΔcyt^Δ/Δ^* mice displayed previously reported defects in axon pathfinding in the anterior commissure, the piriform cortex, lateral olfactory tract, thalamocortical axon tract and corticospinal tract^12,15,17^. Also, neuron migration defects in the cortex and cerebellum were similar in *Sema6a^-/-^* and *Sema6aΔcyt^Δ/Δ^* mice^13,17,25^. Interestingly, knockout of both *Plxna2* and *Plxna4* leads to migration defects in the cerebral cortex similar to those observed in *Sema6a^-/-^* and *Sema6aΔcyt^Δ/Δ^* mice^25^. This suggests that SEMA6A *reverse* signaling in the cortex *in vivo* may be induced by Plexin-A2 and -A4.

Interestingly, dentate gyrus neurons in the hippocampus migrate correctly in the absence of SEMA6A *reverse* signaling (in *Sema6aΔcyt^Δ/Δ^* mice). This indicates that in *Sema6aΔcyt^Δ/Δ^* mice the extracellular part of SEMA6A is intact and functional. Further, these data suggest that the formation of the dentate gyrus infrapyramidal blade relies on SEMA6A *forward* signaling. However, the molecular pathways that act downstream of SEMA6A *forward* signaling during dentate gyrus formation remain unclear. Although severe dentate gyrus malformations are observed in both *Plxna2* and *Plxna4* knockout mice^36^, the observed phenotypes differ from the migration defect found in *Sema6a* mutants. This suggests that other receptors may be involved in mediating the effects of SEMA6A in this case.

In both *Sema6a* knockout and *Plxna2;Plxna4* double knockout mice, TCA bundles aberrantly extend into the ventral telencephalon^12,16^. Intriguingly, thalamic axons express SEMA6A, and SEMA6A, Plexin-A2 and -A4 are co-expressed in guidepost cells along the TCA trajectory. A subset of these guidepost cells is misplaced in both *Sema6a* and *Plxna2;Plxna4* double knockout models^22^. These data hinted at a role for SEMA6A *reverse* signaling in TCAs. Our *Sema6aΔcyt^Δ/Δ^* mice confirm a cell-autonomous role for SEMA6A as a receptor on TCAs responsible for their correct pathfinding. Whether SEMA6A in guidepost cells has additional roles in TCA pathfinding remains to be shown.

In this study, we compared a select number of axonal and neuronal phenotypes in *Sema6a^-/-^* and *Sema6aΔcyt^Δ/Δ^* mice. However, *Sema6a* is broadly expressed in the developing nervous system. Further work is needed to address whether the SEMA6A *reverse* signaling pathway also operates in other regions, such as the retina or spinal cord. The ability to conditionally remove the SEMA6A intracellular domain in *Sema6aΔcyt^fl/fl^* mice *in vivo* opens up new possibilities to investigate the receptor functions of SEMA6A, both at a cellular and molecular level.

## METHODS

### Construct design

Our constructs are designed based on *Sema6a-transcript 001* (6856 bp, 1031aa, ENSMUST00000019791). Full-length *Sema6a* cDNA or shortened *Sema6a* lacking the sequence coding for the intracellular domain (bp 2680-3766) were cloned into the *pCAG-GFP* vector (Addgene, Cat #11150) to obtain *pCAG*-promotor-driven expression of *Sema6a* and *Sema6aΔcyt* with a GFP signal sequence located upstream.

### Cell culture

To validate protein expression and membrane localization *pCAG-Sema6a-GFP* or *pCAG-Sema6aΔcyt-GFP* constructs were transfected in N2A cells. To assess whether the *Sema6aΔcyt* construct produced a truncated SEMA6A protein able to elicit forward signaling as ligand for Plexin-As, all three constructs were studied in NIH3T3-cortical explant co-cultures. Cell culture methods were as previously described^37^. In brief, N2A or NIH3T3 cells were plated 2000 cells/m and cultured in DMEM high glucose (GIBCO) containing 10% fetal calf serum, L-glutamine and penicillin/streptomycin. At the day of transfection, when the cells reached 70-80% confluency, they were switched to DMEM high glucose medium without antibiotics 1 h prior to transfection. Cells were transfected with 1 µg of *pCAG-GFP* (only in NIH3T3), *pCAG-Sema6a-GFP* or *pCAG-Sema6aΔcyt-GFP* constructs in 1:3 PEI 1 mg/ml (Polyscience) in H2O, supplemented with 150 mM NaCl. The next day, the transfection medium was replaced by DMEM high glucose without antibiotics.

Primary cortical explants were generated from P0-P4 *wildtype* mice. Pups were decapitated, brains were rapidly removed, kept in ice-cold L15 dissection medium and 350 μm-thick brain slices were made with a tissue chopper (McIllwain). The cortex was dissected and cut into small pieces of 100-300 μm. Explants were plated on NIH3T3 cells expressing either *pCAG-GFP* control, or *pCAG-Sema6a-GFP* or *pCAG-Sema6aΔcyt-GFP* constructs. N2A cells and NIH3T3-cortical explant co-cultures were grown for 48 and 72 h respectively, and fixed by adding equal volume of 8% PFA in PBS containing 30% sucrose for 10-30 min at room temperature followed by PBS wash and immunocytochemistry analysis.

### Immunocytochemistry

All antibodies used in this study are listed in table 1. For surface expression analysis, N2A cells were stained by adding goat-anti-SEMA6A antibodies (1:1000, R&D, AF1615) to the culture medium for 10 minutes at 37 °C, 5% CO2. Cells were washed and fixed by adding 1:1 8% PFA supplemented with 30% sucrose in PBS to the medium for 1 h at 37 °C. After PBS washes, cells were incubated in blocking buffer (PBS with 5% BSA) for 30 min and incubated with Alexa Fluor-568-conjugated donkey anti goat secondary antibodies (1:1000, Invitrogen) in PBS with 2.5% BSA for 1 h at RT. Subsequently, cells were treated following a triton-based immunocytochemistry staining protocol. N2A or NIH3T3-cortical explant co-cultures were fixed with 4% PFA in PBS for 10-30 minutes at room temperature and permeabilised with 0.1% triton in 5% BSA-PBS blocking buffer for 30 min at RT. The cytosolic pool of SEMA6A-GFP or SEMA6AΔcyt-GFP was visualised using rabbit-anti-GFP (1:1000, A11122 Life Technologies) followed by 1 h incubation of Alexa Fluor-488-conjugated donkey-anti-rabbit secondary antibodies (1:1000) in PBS. Neurite outgrowth of cortical explants was visualized using mouse-anti-Tuj1 antibodies (1:500, MMS-435P Covance) combined with Alexa Fluor-647-conjugated donkey-anti-mouse secondary antibodies (1:1000, Invitrogen). Counterstaining with Hoechst 33258 (#10778843 Fisher) was used to visualize cell nuclei. Coverslips were mounted using Fluorsave. Images were taken on an Axioskop EPI-fluorescent microscope (Zeiss) and processed using FIJI image analysis. Surface SEMA6A expression in N2A cells was quantified by analyzing the mean intensity of GFP signal divided by the mean intensity of SEMA6A staining per cell. Neurite outgrowth of cortical explants was quantified by calculating a ratio between the total area of the explant including neurites, with the area of the explant core. Data were plotted and statistical analysis was performed using GraphPad Prism 9.

**Table 1.** Antibody list.

| **Antibody** | **Dilution** | **Reference #** | **RRID** |
| --- | --- | --- | --- |
| *Primary antibodies* |  |  |  |
| NeuN (mouse) | 1:000 | Abcam, Cat# ab104224 | AB_10711040 |
| Prox1 (rabbit) | 1:1000 | Abcam, Cat# ab101851 | AB_10712211 |
| L1CAM (rat) | 1:500 | Millipore, Cat# MAB5272 | AB_2133200 |
| BrdU (rat) | 1:500 | Accurate, Cat# OBT0030 | AB_2313756 |
| SEMA6A (goat) | 1:1000 | R&D, Cat # AF1615 | AB_2185995 |
| Tuj1 (mouse) | 1:500 | Covance, Cat# MMS-435P | AB_2313773 |
| GFP (rabbit) | 1:1000 | Life Technologies, Cat# A11122 | AB_221569 |
| Tbr1 (rabbit) | 1:200 | Abcam, Cat# ab31940 | AB_2200219 |
| Ctip2 (rat) | 1:1000 | Abcam, Cat# ab18465 | AB_2064130 |
| Satb2 (rabbit) | 1:500 | Abcam, Cat# ab34735 | AB_2301417 |
| Reelin (mouse) | 1:1000 | MBL International Corporation, Cat# D223-3 | AB_843523 |
| *Secondary antibodies* |  |  |  |
| Donkey-anti-mouse 488 | 1:750 (IHC) | Thermo Fisher Scientific, Cat# A21202 | AB_141607 |
| Donkey-anti-rabbit Alexa 488 | 1:750 (IHC) | Thermo Fisher Scientific, Cat# A21206 | AB_2535792 |
| Donkey-anti-goat Alexa 568 | 1:750 (IHC) | Abcam, Cat# ab175474 | AB_2636995 |
| Donkey-anti-rabbit Alexa 568 | 1:750 (IHC) | Abcam, Cat# ab175470 | AB_2783823 |
| Donkey-anti-rat Alexa 568 | 1:750 (IHC) | Thermo Fisher Scientific, Cat# A78946 | AB_2910653 |
| Donkey-anti-rat Alexa 647 | 1:750 (IHC) | Abcam, Cat# ab150155 | AB_2813835 |
| Donkey-anti-mouse Alexa 647 | 1:750 (IHC) | Thermo Fisher Scientific, Cat# A31571 | AB_162542 |
| Donkey-anti-mouse Alexa IR800 | 1:10000 (WB) | LI-COR Biosciences, Cat# 926-32212 | AB_621847 |
| Donkey-anti-goat Alexa 680 | 1:10000 (WB) | Jackson ImmunoResearch, Cat# 705-625-147 | AB_2340440 |
| Donkey-anti-goat Alexa 680 | 1:10000 (WB) | Jackson ImmunoResearch, Cat# 705-625-147 | AB_2340440 |
| Donkey-anti-goat HRP | 1:20000 (WB) | Abcam, Cat# ab205723 | AB_3065024 |
| Goat-anti-rabbit HRP | 1:20000 (WB) | Abcam, Cat# ab205718 | AB_2819160 |

### Animals

All animal use and care were in accordance with institutional guidelines and approved by the Animal Ethics Committee of Utrecht University (Dierexperimenten Ethische Commissie) (CCD licence: AVD115002016532) and conducted in agreement with Dutch laws (Wet op de Dierproeven, 1996, revised 2014) and European regulations.

To assemble the *Sema6aΔcyt* mouse cassette, *Sema6a* full-length exon 19 and a neomycin cassette surrounded by *Frt* sites, was flanked with *LoxP* sites. A shortened version of exon 19, lacking the coding sequence for the cytoplasmic domain but including the 3’ untranslated region, was placed behind it. This combination, flanked with homologous targeting sequences was cloned into pcDNA3.1 (Invitrogen, Cat #V79020). The targeting vector was electroporated in ES cells at Biocytogen. Homologous recombination was assessed using southern blot (data not shown). The integration of a *LoxP* site in the large sequence of intron 18 did not interfere with mRNA splicing. *Sema6aΔcyt^fl/wt^* F1 founders were generated at Biocytogen, and the neomycin cassette was removed by *Flp* recombination. *Cre* recombination of Sema6a*Δcyt^fl^* mice results in the removal of the cassette containing the full-length exon 19, which is replaced by a mutated exon 19 sequence encoding only the transmembrane domain of SEMA6A (Fig. 1G).

The *Sema6a* mouse strains used in this study, *Sema6aΔcyt^fl/wt^*and *Sema6a^-/-^* (Sema6aGt[KST069]Byg)^12^ (kindly provided by Kevin Mitchell) were kept on a *C57Bl/6J* background. *C57Bl6/J wildtype* mice were used as controls, unless otherwise specified. Timed-pregnant females were 3-6 months of age. The morning on which a vaginal plug was detected was considered embryonic day 0.5 (E0.5). We crossed *Sema6aΔcyt* mice (F1 created at Biocytogen) with specific *Cre* lines. To obtain *Cre*-recombination in the cortex and hippocampus specifically, *Sema6aΔcyt* mutants were mated with and *Emx1^IRES^ ^Cre^* (JAX stock #005628)^33^ resulting in *Emx1^Cre^;Sema6aΔcyt* mice. To delete SEMA6A’s intracellular domain from the germline, *Sema6aΔcyt* mutants were mated with *EIIa-Cre* (JAX stock #003724)^32^, resulting in *Sema6aΔcyt^Δ/Δ^* animals (see table 2 for strain details). To identify *LoxP* sites and check for germline recombination we obtained tail samples for DNA extraction and PCR analysis was done using primers and protocols as described in table 3 (PCR results can be seen in Fig. 1H,I).

**Table 2.** Mouse lines.

| **Mouseline** | **Reference** | **Source** | **RRID** |
| --- | --- | --- | --- |
| *Ella-Cre* | The Jackson Laboratory <sup>32</sup> | JAX:003724 | IMSR_JAX:003724 |
| *EMX1-IRES-Cre (Emx1<sup>tm1Cre</sup>)<sup>Krj</sup>* | The Jackson Laboratory <sup>33</sup> | JAX:005628 | IMSR_JAX:005628 |
| *Sema6a<sup>-/-</sup> (Sema6aGt[KST069]Byg)* | <sup>12</sup> | Provided by<br>Kevin Mitchell | MGI:3037891 |
| *Sema6aΔcyt<sup>fl/+</sup>* | Biocytogen |  |  |
| *C57BL/6J* | Charles Rivers Laboratories |  | IMSR_JAX:000664 |

**Table 3.**
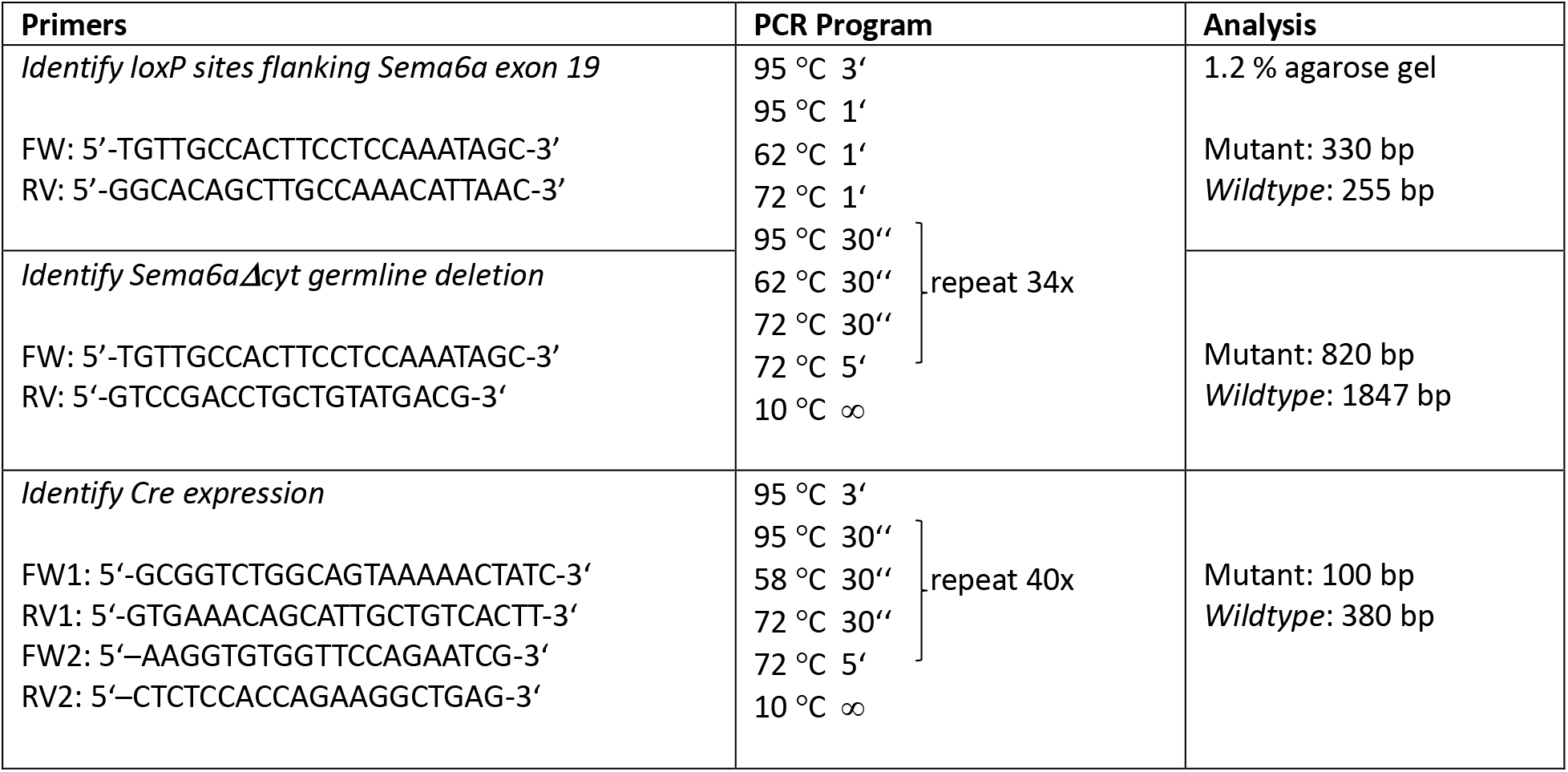
*Sema6aΔcyt* mouse genotyping procedure.

### SEMA6AΔcyt protein expression and localization

To validate *wildtype* versus truncated protein expression, N2As were transfected with *pCAG-Sema6a-GFP* and *pCAG-Sema6aΔcyt-GFP* constructs as described above. 48 hours after transfection, N2As were lysed with RIPA buffer (10mM Tris-buffer pH 7.5, 150 mM NaCl, 0.1% SDS, 1% Triton X-100, 1% Deoxycholate, 0.5 mM EDTA, 1 mM PMSF and complete protease inhibitor (Roche) and spun down at 12.000 RPM at 4°C for 15 min. The protein containing supernatant was isolated and transferred to a new tube.

To asses SEMA6A localisation to the membrane in *Emx1^Cre^;Sema6aΔcyt^fl/fl^*and *Emx1^Cre^;Sema6aΔcyt^fl/wt^* control mice, membrane fractions where prepared using a sucrose buffer (adapted protocol^38^). In short, brain tissue was dissected into two samples containing cortices and hippocampi (sample 1) separated from the rest of the brain (sample 2) and snap-frozen in dry ice. Three volumes of fractionation buffer were added to the tissue (0.25 M sucrose, 1 mM EDTA, 5 mM NaCl, 10 mM KCl, protease inhibitor 1x and 10 mM Tris-HCl pH7.5) followed by homogenization. Nuclei were removed by centrifugation at 1000 g (5000rpm) for 6 min at 4 °C. The supernatant was centrifuged at 9.200 g (12500 rpm) at 4 °C for 15 min to obtain cytosolic and membrane fractions. These fractions were separated by centrifugation at 100.000 g at 4 °C (41000 rpm TLA 45 rotor) for 2 h, resulting in a cytosol enriched supernatant and a plasma membrane enriched pellet. The cytosol was discarded and the membrane pellet was washed in sucrose buffer and centrifugated at 100.000 g at 4 °C for 1 h. The pellet was resuspended in sucrose buffer.

For western blot analysis, protein samples were supplemented with sample buffer (NuPage, 1:4) and 10% B-mercaptoethanol. Samples were denatured for 10 min at 70 °C and loaded (50 µg) on an 8% gel for SDS-polyacrylamide gel electrophoresis. Protein was transferred from gel to a nitrocellulose membrane (Bio-Rad). For N2A lysates, blots were incubated with the following antibodies: goat-anti-SEMA6A (1:1000, AF1615 R&D), rabbit-anti-GFP (1:1000, Life Technologies, A11122) followed by the appropriate HRP-conjugated secondary antibody in 1.5% milk in TBS-T. Immunoblots were exposed to Pierce ECL substrate (Thermo Fisher Scientific, 32106) and digitized using an Epson flatbed scanner (Perfection 4990, Epson America). For mouse tissue membrane fractions, blots were incubated with goat-anti-SEMA6A (1:1000, AF1615 R&D) and mouse-anti-Tuj1 (1:500, MMS-435P Covance) followed by fluorescent secondary antibodies donkey-anti-mouse Alexa-IR800 (1:10000) and donkey-anti-goat Alexa-680 (1:10000, Invitrogen). Fluorescence was detected using the Odyssey CLx imaging system (Licor).

### BrdU injections

To assess granule cell proliferation and migration patterns in the cerebellum we used 5-Bromo-2’-deoxyuridine (BrdU) in *Sema6a^-/-^*, *Sema6aΔcyt^Δ/Δ^* and *wildtype* mice. P15 pups were injected intraperitoneally with BrdU (Sigma-Aldrich, 15 mg/ml) 50 mg/kg^-1^ in sterile saline solution and placed back in the cage with their litter mates and mother. At P30, animals were killed by intraperitoneal injection of an overdose of pentobarbital (Euthanimal, Alfasan) and transcardially perfused with 4% PFA in PBS. Brains were post-fixed in 4% PFA overnight, cryoprotected in 30% sucrose in PBS, and snap-frozen in -40 °C isopentane. 20 µm sagittal cryosections were made using a Leica CM3050 cyrostat and processed for immunohistochemistry.

### Immunohistochemistry

Free-floating brain sections were washed with PBS. Tissue was incubated in blocking buffer (PBS with 5% BSA and 0.1% triton-X-100) for 1 h at RT. Primary antibodies (as specified in results section and listed in table 1) were incubated overnight at 4 °C in 2.5% BSA and 0.1% triton-X in PBS. Sections were washed with PBS and incubated with secondary antibodies for 2 h at RT in 2.5% BSA and 0.1% triton-X in PBS followed by counterstaining with 4’,6-diamidino-2-fenylindool (DAPI) (1µg/µl, Sigma, D9564) for 15 minutes at RT. To assess neuroanatomical phenotypes, at least 3 animals per genotype were analyzed.

For BrdU immunohistochemistry antigen retrieval was required. Sections were incubated in 2 N HCl for 30 min at 65 °C followed by 10 min incubation in 0.1 M boric acid, pH 8.0. Sections were washed with PBS and further processed following blocking and antibody incubation as described above, using rat-anti-BrdU (1:500, OBT0030 Accurate) and donkey-anti-rat Alexa Fluor-568-conjugated secondary antibody (1:1000, Invitrogen).

Brain sections were mounted using Fluorsave, imaged on an Axioskop EPI-fluorescent microscope (Zeiss) and processed using FIJI image analysis. To assess BrdU labeling in the cerebellum, the molecular layer and internal granule cell layer were defined as ROIs. The BrdU channel was binarized, followed by particle analysis using the partical analysis plugin of FIJI. The number of BrdU-positive cells was divided by the area of the ROI to calculate the number of cells/µm^2^. To analyze the proportion of cells that had migrated properly into the internal granule layer, we calculated the ratio between cells/um^2^ in the internal granule layer and molecular layer. We counted and averaged cells from at least 10 sections of 7 control brains (including 2 *wildtype*, 2 *Sema6a^+/-^* and 3 *Sema6aΔcyt^Δ/wt^*), 2 *Sema6a^-^/-* and 3 *Sema6aΔcyt^Δ/Δ^*. Data were plotted and statistical analysis was performed using Graphpad Prism 9 software.

## ACKNOWLEDGEMENTS

We thank all members of the Pasterkamp laboratory and Department of Translational Neuroscience for assistance and helpful discussions throughout this project. We thank Kevin Mitchell for kindly providing *Sema6a^-/-^* mice, and Biocytogen for generating *Sema6aΔcyt* animals. This work was supported by the Netherlands Organization for Health Research and Development (ALW-VICI). R.J.P. is funded through the Gravitation program of the Dutch Ministry of Education, Culture, and Science and the Netherlands Organization for Scientific Research (BRAINSCAPES).

## AUTHOR CONTRIBUTIONS

M.G.V., S.L. and R.J.P. designed all experiments. M.G.V., K.J., E.Y.vB. and M.H.vdM. performed *in vivo* experiments. S.L., A.K. and M.G.V. performed *in vitro* experiments. C.M. and N.H.C.vK. arranged mouse breeding. S.L., K.R., Y.A. and M.G.V. generated *Sema6a* constructs. M.G.V., E.Y.vB., M.H.vdM. and R.J.P. wrote the manuscript with input from all authors. R.J.P. acquired funding, conceptualized the project and provided supervision.

## REFERENCES

1. Goodman CS. MECHANISMS AND MOLECULES THAT CONTROL I chemoattraction I chemorepulsion I. Annual Reviews Neuroscience. Published online 1996.

2. Tessier-Lavigne M, Goodman CS. The Molecular Biology of Axon Guidance. Science (1979). 1996;274(5290):1123-1133. doi:10.1126/science.274.5290.1123

3. Tamagnone L, Artigiani S, Chen H, et al. Plexins are a large family of receptors for transmembrane, secreted, and GPI-anchored semaphorins in vertebrates. Cell. 1999;99(1):71–80. doi:10.1016/S0092-8674(00)80063-X

4. Pasterkamp RJ. Getting neural circuits into shape with semaphorins. Nat Rev Neurosci. 2012;13(9):605–618. doi:10.1038/nrn3302

5. Robinson RA, Griffiths SC, van de Haar LL, et al. Simultaneous binding of Guidance Cues NET1 and RGM blocks extracellular NEO1 signaling. Cell. 2021;184(8):2103–2120.e31. doi:10.1016/j.cell.2021.02.045

6. Jongbloets BC, Pasterkamp RJ. Semaphorin signalling during development. Development. 2014;141(17):3292–3297. doi:10.1242/dev.105544

7. Zang Y, Chaudhari K, Bashaw GJ. New insights into the molecular mechanisms of axon guidance receptor regulation and signaling. Curr Top Dev Biol. 2021;142:147–196. doi:10.1016/bs.ctdb.2020.11.008

8. Renaud J, Kerjan G, Sumita I, et al. Plexin-A2 and its ligand, Sema6A, control nucleus-centrosome coupling in migrating granule cells. Nat Neurosci. 2008;11(4):440-449. doi:10.1038/nn2064

9. Haklai-Topper L, Mlechkovich G, Savariego D, Gokhman I, Yaron A. Cis interaction between Semaphorin6A and Plexin-A4 modulates the repulsive response to Sema6A. EMBO J. 2010;29:2635–2645. doi:10.1038/emboj.2010.147

10. Matsuoka RL, Nguyen-Ba-Charvet KT, Parray A, Badea TC, Chédotal A, Kolodkin AL. Transmembrane semaphorin signalling controls laminar stratification in the mammalian retina. Nature. 2011;470(7333):259-264. doi:10.1038/nature09675

11. Perez-Branguli F, Zagar Y, Shanley DK, Graef IA, Chédotal A, Mitchell KJ. Reverse signaling by semaphorin-6A regulates cellular aggregation and neuronal morphology. PLoS One. 2016;11(7):1–24. doi:10.1371/journal.pone.0158686

12. Leighton PA, Mitchell KJ, Goodrich L V, et al. Defining brain wiring patterns and mechanisms through gene trapping in mice. Nature. 2001;410(March):1–6. papers2://publication/uuid/F14E2A4D-06C2-4EAA-9E4D-E6A9B33EE66D

13. Kerjan G, Dolan J, Haumaitre C, et al. The transmembrane semaphorin Sema6A controls cerebellar granule cell migration. Nat Neurosci. 2005;8(11):1516–1524. doi:10.1038/nn1555

14. Suto F, Tsuboi M, Kamiya H, et al. Interactions between Plexin-A2, Plexin-A4, and Semaphorin 6A Control Lamina-Restricted Projection of Hippocampal Mossy Fibers. Neuron. 2007;53(4):535-547. doi:10.1016/j.neuron.2007.01.028

15. Rünker AE, Little GE, Suto F, Fujisawa H, Mitchell KJ. Semaphorin-6A controls guidance of corticospinal tract axons at multiple choice points. Neural Dev. 2008;3(1). doi:10.1186/1749-8104-3-34

16. Little GE, López-Bendito G, Rünker AE, et al. Specificity and plasticity of thalamocortical connections in Sema6A mutant mice. PLoS Biol. 2009;7(4):0756–0770. doi:10.1371/journal.pbio.1000098

17. Rünker AE, O’Tuathaigh C, Dunleavy M, et al. Mutation of Semaphorin-6A disrupts limbic and cortical connectivity and models neurodevelopmental psychopathology. PLoS One. 2011;6(11). doi:10.1371/journal.pone.0026488

18. Sun LO, Jiang Z, Rivlin-Etzion M, et al. On and Off Retinal Circuit Assembly by Divergent Molecular Mechanisms. Science (1979). 2013;342(6158):1241974-1241974. doi:10.1126/science.1241974

19. Renaud J, Chédotal A. Time-lapse analysis of tangential migration in Sema6A and PlexinA2 knockouts. Molecular and Cellular Neuroscience. 2014;63:49–59. doi:10.1016/j.mcn.2014.09.005

20. Sun LO, Brady CM, Cahill H, et al. Functional Assembly of Accessory Optic System Circuitry Critical for Compensatory Eye Movements. Neuron. 2015;86(4):971–984. doi:10.1016/j.neuron.2015.03.064

21. Belle M, Parray A, Belle M, Chédotal A, Nguyen-Ba-Charvet KT. PlexinA2 and Sema6A are required for retinal progenitor cell migration. Dev Growth Differ. 2016;58(5):492–502. doi:10.1111/dgd.12298

22. Mitsogiannis MD, Little GE, Mitchell KJ. Semaphorin-Plexin signaling influences early ventral telencephalic development and thalamocortical axon guidance. Neural Dev. 2017;12(1):1–20. doi:10.1186/s13064-017-0083-4

23. Okada T, Keino-masu K, Suto F, Mitchell KJ, Masu M. Remarkable complexity and variability of corticospinal tract defects in adult Semaphorin 6A knockout mice. Brain Res. 2019;1710(June 2018):209–219. doi:10.1016/j.brainres.2018.12.041

24. Lilley BN, Sabbah S, Hunyara JL, et al. Genetic access to neurons in the accessory optic system reveals a role for Sema6A in midbrain circuitry mediating motion perception. Journal of Comparative Neurology. 2019;527(1):282–296. doi:10.1002/cne.24507

25. Hatanaka Y, Kawasaki T, Abe T, et al. Semaphorin 6A–Plexin A2/A4 Interactions with Radial Glia Regulate Migration Termination of Superficial Layer Cortical Neurons. iScience. 2019;21:359–374. doi:10.1016/j.isci.2019.10.034

26. Alto LT, Terman JR. Semaphorins and their signaling mechanisms. Methods in Molecular Biology. Published online 2017. doi:10.1007/978-1-4939-6448-2_1

27. Godenschwege TA, Hu H, Shan-Crofts X, Goodman CS, Murphey RK. Bi-directional signaling by semaphorin 1a during central synapse formation in Drosophila. Nat Neurosci. 2002;5(12):1294–1301. doi:10.1038/nn976

28. Yu L, Zhou Y, Cheng S, Rao Y. Plexin A-Semaphorin-1a reverse signaling regulates photoreceptor axon guidance in Drosophila. Journal of Neuroscience. 2010;30(36):12151–12156. doi:10.1523/JNEUROSCI.1494-10.2010

29. Jeong S, Juhaszova K, Kolodkin AL. The Control of Semaphorin-1a-Mediated Reverse Signaling by Opposing Pebble and RhoGAPp190 Functions in Drosophila. Neuron. 2012;76(4):721–734. doi:10.1016/j.neuron.2012.09.018

30. Jeong S, Yang D som, Hong YG, Mitchell SP, Brown MP, Kolodkin AL. Varicose and cheerio collaborate with pebble to mediate semaphorin-1a reverse signaling in Drosophila. Proc Natl Acad Sci U S A. 2017;114(39):E8254–E8263. doi:10.1073/pnas.1713010114

31. Mauti O, Domanitskaya E, Andermatt I, Sadhu R, Stoeckli ET. Semaphorin6A acts as a gate keeper between the central and the peripheral nervous system. Neural Dev. 2007;2(1):28. doi:10.1186/1749-8104-2-28

32. Lakso M, Pichel JG, Gorman JR, et al. Efficient in vivo manipulation of mouse genomic sequences at the zygote stage. Proceedings of the National Academy of Sciences. 1996;93(12):5860–5865. doi:10.1073/pnas.93.12.5860

33. Gorski JA, Talley T, Qiu M, Puelles L, Rubenstein JLR, Jones KR. Cortical excitatory neurons and glia, but not GABAergic neurons, are produced in the Emx1-expressing lineage. Journal of Neuroscience. 2002;22(15):6309–6314. doi:10.1523/jneurosci.22-15-06309.2002

34. Shammah-Lagnado SJ, Alheid GF, Heimer L. Afferent connections of the interstitial nucleus of the posterior limb of the anterior commissure and adjacent amygdalostriatal transition area in the rat. Neuroscience. 1999;94(4):1097–1123. doi:10.1016/S0306-4522(99)90280-4

35. Ma J, Yao XH, Fu Y, Yu YC. Development of layer 1 neurons in the mouse neocortex. Cerebral Cortex. 2014;24(10):2604–2618. doi:10.1093/cercor/bht114

36. Zhao XF, Kohen R, Parent R, et al. PlexinA2 Forward Signaling through Rap1 GTPases Regulates Dentate Gyrus Development and Schizophrenia-like Behaviors. Cell Rep. 2018;22(2):456–470. doi:10.1016/j.celrep.2017.12.044

37. Van Battum EY, Gunput RAF, Lemstra S, et al. The intracellular redox protein MICAL-1 regulates the development of hippocampal mossy fibre connections. Nat Commun. 2014;5(1):4317. doi:10.1038/ncomms5317

38. Van Erp S, Van den Heuvel DMA, Fujita Y, et al. Lrig2 Negatively Regulates Ectodomain Shedding of Axon Guidance Receptors by ADAM Proteases. Dev Cell. 2015;35(5):537–552. doi:10.1016/j.devcel.2015.11.008

